# Human papilloma virus E6 regulates therapy responses in oropharyngeal cancer by repressing the PGC-1α/ERRα axis

**DOI:** 10.1101/2022.05.04.490169

**Authors:** Malay K. Sannigrahi, Pavithra Rajagopalan, Ling Lai, Xinyi Liu, Varun Sahu, Hiroshi Nakagawa, Jalal Jalaly, Robert M. Brody, Iain M. Morgan, Bradford E. Windle, Xiaowei Wang, Phyllis A. Gimotty, Daniel P. Kelly, Elizabeth A. White, Devraj Basu

**Author notes:** **Corresponding author**: Devraj Basu, MD, PhD, 3400 Spruce Street, 5 Ravdin/Silverstein, Philadelphia, PA 19104.

## Abstract

Therapy with radiation plus cisplatin kills human papilloma virus-related (HPV+) oropharyngeal squamous cell carcinomas (OPSCCs) by increasing reactive oxygen species beyond cellular antioxidant capacity. To explore why some patients fail these standard treatments, we evaluated whether the variation in HPV oncoprotein levels among HPV+ OPSCCs impacts mitochondrial metabolism, a source of antioxidant capacity. In cell line and patient-derived xenograft models, levels of HPV full-length E6 (fl-E6) inversely correlated with oxidative phosphorylation, antioxidant capacity, and therapy resistance, and fl-E6 was the only HPV oncoprotein to display such correlation. Ectopically expressing fl-E6 in models with low levels reduced mitochondrial mass, depleted antioxidant capacity, and sensitized to therapy. In this setting, fl-E6 repressed the PGC-1α/ERRα pathway for mitochondrial biogenesis by reducing p53-dependent PGC-1α transcription. Concordant observations were made in three clinical cohorts, where expression of mitochondrial components was higher in tumors of patients with reduced survival. These tumors contained the lowest fl-E6 levels, highest p53 target gene expression, and an activated PGC-1α/ERRα pathway. Our findings demonstrate that E6 can potentiate treatment responses by depleting mitochondrial antioxidant capacity and provide evidence for low E6 negatively impacting patient survival. E6’s interaction with the PGC-1α/ERRα axis has implications for predicting and targeting treatment resistance in OPSCC.

## INTRODUCTION

Human papilloma virus-related (HPV+) cancers comprise over 5% of malignancies worldwide, and rising incidence of HPV+ oropharyngeal squamous cell carcinoma (OPSCC) has made it the most common HPV-related cancer in the US (1). HPV+ OPSCCs are typically sensitive to the combination of radiation plus cisplatin, which is often curative for this disease (2). Impaired DNA repair mechanisms help make HPV+ OPSCCs sensitive as a group to these therapies (3), which kill tumors by increasing reactive oxygen species (ROS) generation beyond cellular antioxidant capacity, leading to uncontrolled damage to DNA, protein, and lipids (4). High cure rates for these patients have sustained interest in de-escalating cisplatin and radiation therapy, whose toxic sequelae leave lifelong disabilities in survivors (5). However, attempting to supplant cisplatin with anti-EGFR therapy was unsuccessful (6, 7), and radiation de-escalation efforts remain a work in progress. Ability to personalize therapy based on more accurate predictive biomarkers would greatly enhance treatment de-escalation efforts for this disease; however, such innovation remains hindered by limited understanding of the mechanisms underlying divergent responses among HPV+ OPSCCs to cisplatin and radiation.

Variation among HPV+ OPSCCs in ability to neutralize oxidative stress could contribute to differences in treatment response and survival. Mitochondria contain a large portion of a cell’s antioxidant machinery and not only neutralize ROS byproducts of oxidative metabolism but also allow cancers to survive extrinsic stressors such as hypoxia and nutrient deprivation (4). NADPH generation within mitochondria fuels the glutathione peroxidase and peroxiredoxin antioxidant systems, which are also partly intrinsic to mitochondria (8). Mitochondrial biogenesis and related antioxidant capacity are upregulated by oncogenic drivers in the RAS pathway (9) and contribute to the aggressive phenotype of RAS-driven cancers by mitigating oxidative stress (10). Less is known about mitochondrial metabolism’s contribution to poor outcomes among HPV+ OPSCCs, which are primarily driven by viral factors and show greater vulnerability as a group to oxidative damage. However, there is emerging evidence that HPV+ OPSCC progression may be enhanced by upregulation of oxidative metabolism by diverse mechanisms (11, 12).

A direct role for HPV oncoproteins in regulating mitochondrial function has not been demonstrated. Many viruses have evolved to evade the type I interferon-mediated innate immune responses initiated on the mitochondrial membrane through RIG-I-MDA5-MAVS signaling (13). Viruses have developed mechanisms to inhibit these signals, disrupt mitochondrial membrane potential, and/or impair mitochondrial function by altering fission-fusion dynamics (13). HPV16’s E6 and E7 oncoproteins jointly suppress innate immune gene expression (14), but it is unknown whether this phenomenon is mediated through effects upon mitochondria. Any suppressive effects of E6 and E7 on mitochondrial function to serve HPV replication might secondarily limit growth of HPV+ OPSCCs in hostile microenvironments and sensitize to cisplatin and radiation. Under this hypothesis, the variable viral oncoprotein levels among HPV+ OPSCCs are predicted to impact treatment responses.

This study evaluated whether variations in HPV oncoprotein levels among HPV+ OPSCCs contribute to diversity in their responses to cisplatin and radiation. We found that high oxidative metabolic gene expression is associated with low HPV E6 levels and reduced survival, whereas high E6 levels correlate with decreased mitochondrial function and sensitivity to therapy. Genetic manipulation of E6 expression in HPV+ cancer cell lines revealed that E6 can suppress mitochondrial biogenesis by depletion of p53, a transcription factor for expression of transcriptional co-activator peroxisome proliferator-activated receptor gamma co-activator 1α (PGC-1α) (15). PGC-1α acts with its DNA-binding nuclear partner, estrogen-related receptor α (ERRα), as a master transcriptional regulator for mitochondrial biogenesis (16). Our findings uncover clinically relevant variations in E6 levels and PGC-1α/ERRα pathway activation that provide a new framework for personalizing therapy.

## RESULTS

### High oxidative metabolic gene expression is associated with decreased HPV+ OPSCC survival

Three cohorts were identified that offer publicly available RNAseq data from treatment-naïve HPV+ OPSCCs with survival outcomes annotation. In TCGA, RNAseq data is available from 53 OPSCCs that express high-risk HPV transcripts (17, 18). We used this data to assess the relationship between oxidative metabolic gene expression and overall survival (OS). Expression was quantified for the MSig database’s Hallmark_Oxidative_Phosphorylation Gene Set (19), which contains 200 genes involved in oxidative metabolism and/or the TCA cycle. Upregulation for each gene was defined as expression ≥ one standard deviation above the mean log_w_-transformed level for all cases. The number of upregulated transcripts was used to divide the cases into tertiles. The lowest tertile had better 3-year OS than the highest (p=0.019) and intermediate tertiles (p=0.043) (Figure 1, left). This finding was tested in the JHU HPV+ OPSCC cohort (20). Dividing this cohort (n=47) into tertiles by the identical methodology used for TCGA cases (Figure 1, middle) showed better 3-year OS for the lowest vs. the highest tertile (p=0.048). Wide variation in therapy was apparent from characteristics of the JHU and TCGA cohorts (Supplemental Table 1), making the relationship between survival and response to radiation plus chemotherapy unclear. Thus, we examined a sub-cohort of patients with the broader VU HPV+ OPSCC cohort (21) who received only radiation plus cisplatin as primary therapy (n=37). Despite small sample size, dividing these cases into tertiles revealed better 3-year OS (p=0.041) in the lowest vs. the highest tertile (Figure 1, right). This association of high expression of mitochondrial components with decreased survival across three cohorts led to the hypothesis that high mitochondrial mass contributes to the resistance to cisplatin and radiation seen in a minority of HPV+ OPSCCs.

**Figure 1.**
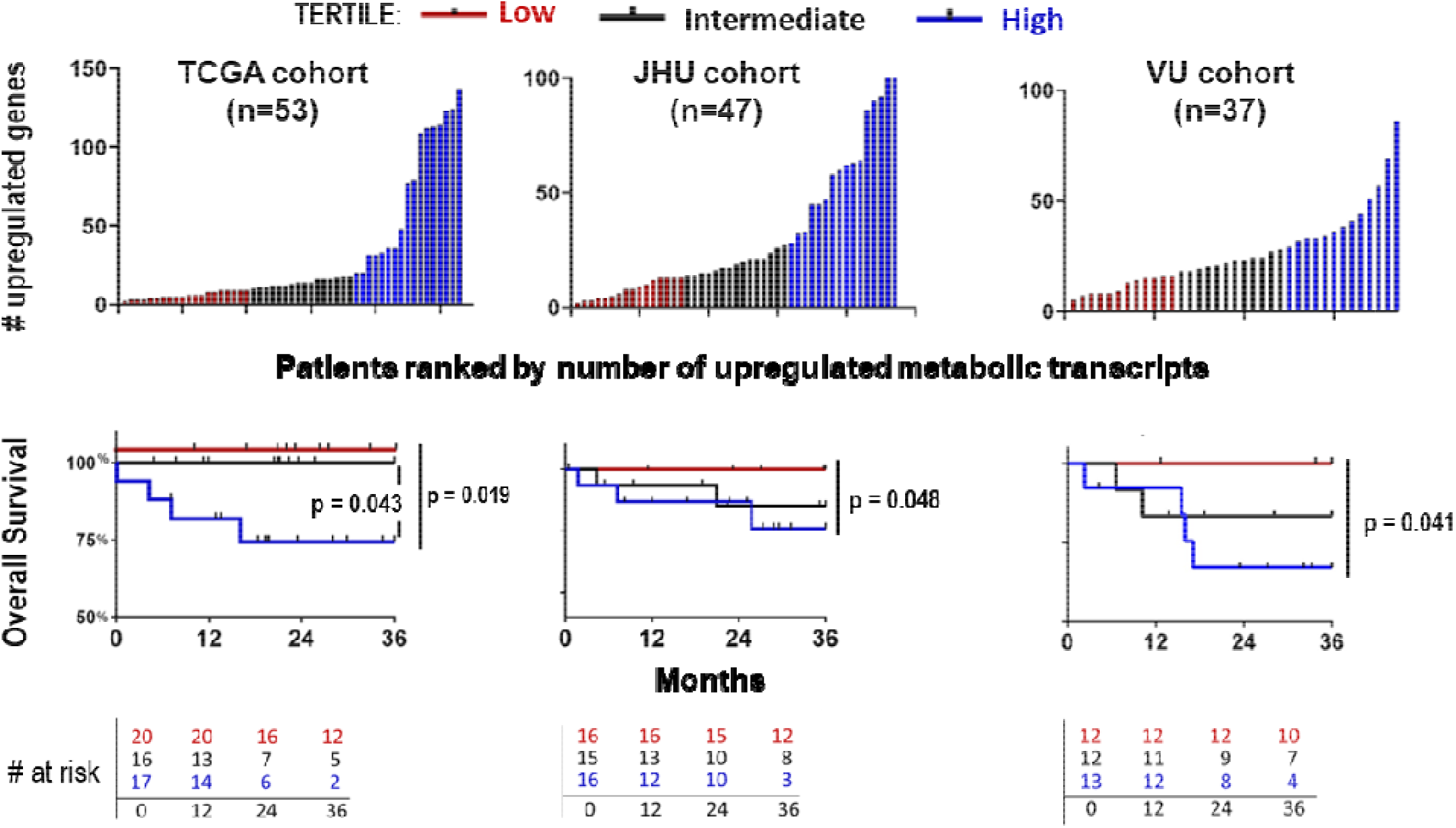
Association of high tumor oxidative metabolic gene expression with reduced patient survival. Three HPV+ OPSCC patient cohorts were divided into tertiles based on the number of upregulated Hallmark_Oxidative_Phosphorylation Gene transcripts in each cohort (top). Upregulation was defined as log_10_-transformed expression of one standard deviation above the mean for each cohort. When a dividing line between tertiles spanned multiple patients with equal upregulated transcripts, those patients were grouped into the higher tertile. Overall survival was estimated by the Kaplan-Meier method (bottom), and the log-rank test was used for pairwise comparisons among tertiles.

### Linking features of poor prognosis patients to high mitochondrial function and cisplatin resistance

The oxidative metabolic gene expression seen in patients with reduced survival was evaluated in patient-derived xenograft (PDX) and cell line models for prediction of increased mitochondrial mass, mitochondrial function, and cisplatin response. RNAseq was performed for the seven HPV+ OPSCC PDX models previously established by us from treatment-naïve patients (22). Up-regulated oxidative metabolic transcripts in the PDXs were quantified by the methodology used in Figure 1. Mitochondrial mass in the PDXs was estimated using levels of mitochondrial DNA measured by a qPCR assay that defines the ratio of a mitochondrial gene *MTCO1* to a nuclear gene *B2M* and then infers mitochondrial DNA levels from a standard curve (23) (Supplemental Figure 1A). A positive linear correlation between the number of upregulated transcripts in the PDXs and *MTCO1/B2M* (Figure 2A, left) supported that our method of quantifying oxidative metabolic gene expression effectively estimates mitochondrial mass. Cisplatin treatment of the PDXs was performed *in vivo* to define the relationship between mitochondrial mass and therapy response. Cisplatin resistance was quantified using rate-based tumor/control (T/C) values, which measures growth of treated tumors relative to controls using an exponential model that incorporates all data points in both growth curves (24) (Supplemental Figure 1B). Rate-based T/C values in the PDXs grown in NOD/SCID/IL-2Rγ^−/−^ (NSG) mice showed a positive linear correlation with mitochondrial mass defined by *MTCO1/B2M* ratio (Figure 2A, right). Relationships among mitochondrial mass, mitochondrial function, and cisplatin resistance were further evaluated in a panel of seven HPV+ cancer cell lines of head and neck origin. Oxidative phosphorylation was quantified by measuring basal oxygen consumption rate (OCR) using the Cell Mito Stress Test on the Seahorse XF™ platform. A strong positive correlation was confirmed between mitochondrial mass and basal OCR in the cell lines (Figure 2B, left, Supplemental Figure 1C). In addition, NADPH/NADP+ ratio was used to quantify antioxidant capacity and showed strong positive correlation with mitochondrial mass (Figure 2B, right). Lastly, cisplatin IC_50_ in the cell lines showed linear positive correlation with *MTCO1/B2M* ratio, basal OCR, and NADPH/NADP+ ratio (Figure 2C). These findings in preclinical models show that the elevated oxidative metabolic gene expression in poor prognosis patients correlates with high mitochondrial mass, oxidative metabolism, and antioxidant capacity. This analysis provided models for subsequent experiments to pursue mechanistic determinants of these metabolic features and assess their impact on therapy response.

**Figure 2.**
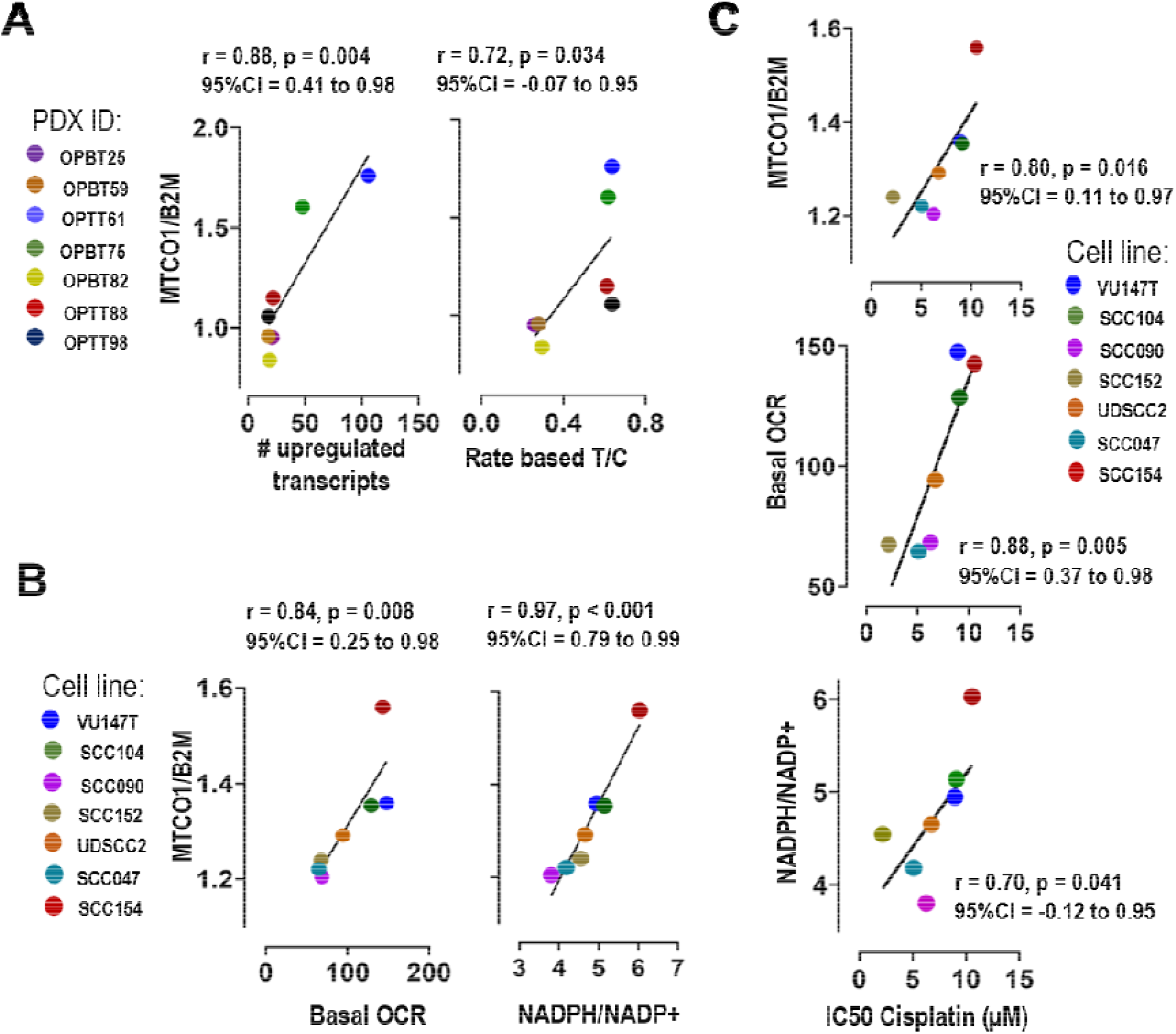
Correlation analyses among oxidative metabolic gene expression, mitochondrial mass, oxidative phosphorylation, and cisplatin resistance in HPV+ OPSCC models. **(A)** Mitochondrial mass was measured in a panel of HPV+ PDXs (n=7) by DNA qPCR to define *MTCO1/B2M*. Correlation analysis was performed for *MTCO1/B2M* vs. # upregulated Hallmark_Oxidative_Phosphorylation genes by RNAseq (left) and vs. cisplatin resistance *in vivo* (right) measured using rate-based T/C value. **(B)** Correlation analysis was performed in cell lines (n=7) for *MTCO1/B2M* vs. basal OCR by Seahorse Assay (left) and vs. NADPH/NAP+ measured by enzyme cycling-based colorimetric assay (right). **(C)** Correlation analysis was performed in cell lines for cisplatin IC_50_ by WST assay vs. *MTCO1/B2M*, basal OCR, and NADPH/NAP+. Pearson correlation coefficients were used to calculate r values for scatter plots with confidence intervals, and p values were determined by t-distribution.

### Among HPV oncogenes, high fl-E6 is uniquely associated with low mitochondrial mass and function

Diversity in viral transcript levels among HPV+ OPSCCs led us to evaluate potential associations between viral gene expression and mitochondrial mass. Among the HPV+ OPSCCs in TCGA, wide variation was apparent among levels of the early viral transcripts with known roles in malignant transformation (E2, E4, E5, E6, E7) (Figure 3A, left). The highest coefficient of variance was seen for the full length, nonspliced form of E6 (fl-E6), a major cancer driver and is the only form of E6 that targets p53. Fl-E6 levels varied by >200-fold (Figure 3A, right) and were reduced in the highest tertile for oxidative metabolic gene expression relative to the lowest tertile (Figure 3B, left). This association was lost when total E6 transcripts or spliced forms (E6*) were considered (Figure 3B, right). No other oncogenic viral transcripts showed association with oxidative metabolic gene expression in TCGA (Supplemental Figure 2A). Although E6 splicing data was not obtainable for the JHU and VU patient cohorts, associations similar to those observed for fl-E6 in TCGA were also present in experimental models of HPV+ OPSCC. In the PDXs, fl-E6 showed negative linear correlation with upregulation of oxidative metabolic transcripts and mitochondrial mass (Figure 3C), whereas no correlations were detected for other oncogenic viral transcripts (Supplemental Figure 2B). Similarly, fl-E6 was negatively correlated with *MTCO1/B2M* ratio, basal OCR, and NADPH/NADP+ in the cell lines (Figure 3D), whereas such relationships were not apparent for other oncogenic HPV transcripts (Supplemental Figure 2C). These findings reveal a reciprocal relationship between viral oncogene expression and both mitochondrial mass and mitochondrial function that is specific to fl-E6.

**Figure 3.**
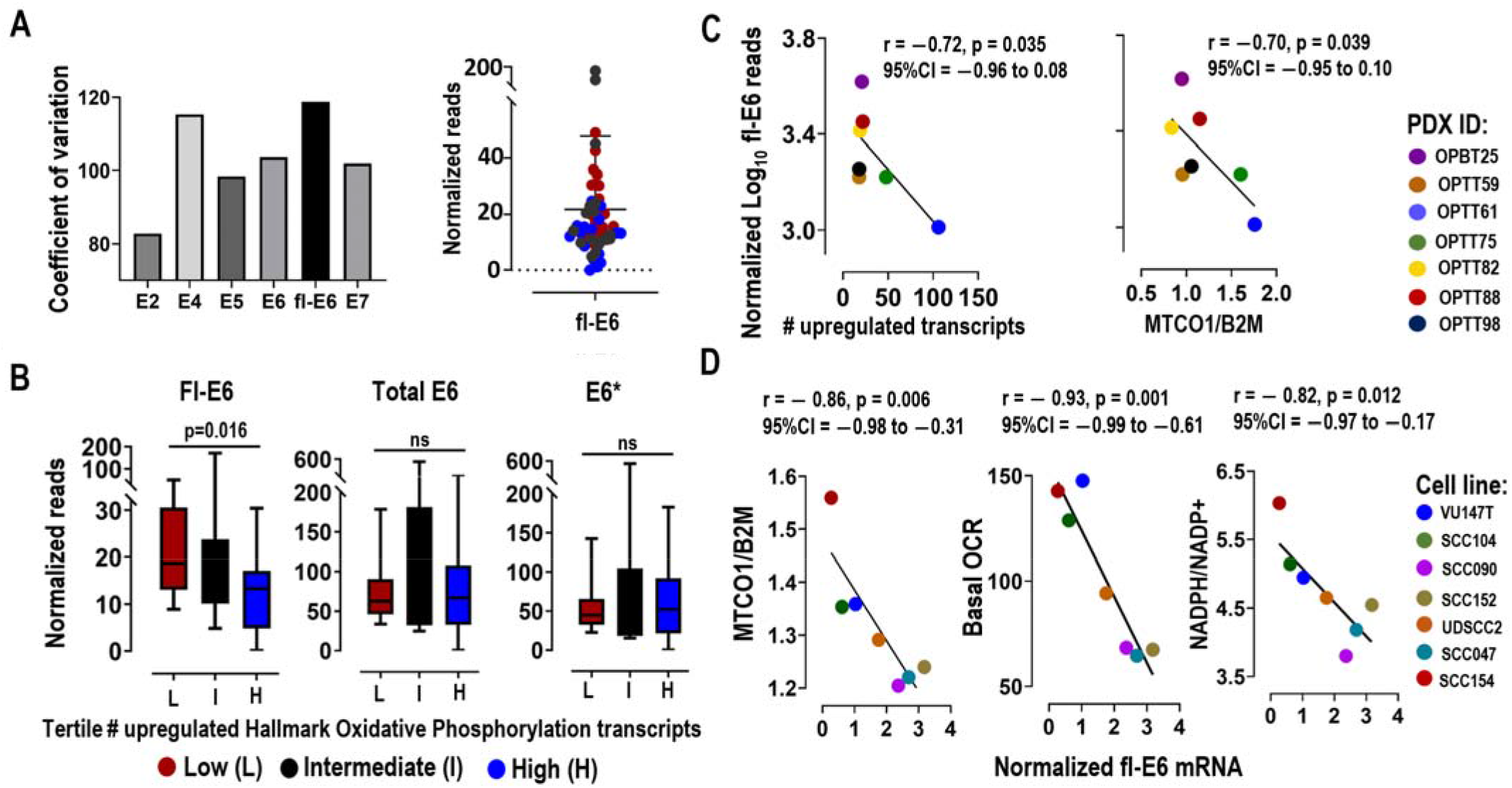
Fl-E6 levels are associated with low mitochondrial mass and function in patient tumors and models. **(A)** RNAseq data for the HPV+ OPSCCs in TCGA (n=53) showing coefficients of variation among the mRNAs for viral oncogenes (left) and the range of fl-E6 expression across the cohort (right). **(B)** RNAseq comparison of fl-E6, total E6, and E6* levels (right) between lowest and highest tertiles for Hallmark_Oxidative_Metabolism gene expression in TCGA. P values were calculated by the Mann Whitney Test. n.s.=not significant. **(C)** PDX fl-E6 mRNA levels by RNAseq vs. number of up-regulated Hallmark_Oxidative_Phosphorylation Genes (left) and vs. mitochondrial mass (right). **(D)** Cell line fl-E6 levels normalized to 18S by qRT-PCR vs. mitochondrial mass (*MTCO1/B2M*), basal OCR (Seahorse Assay), and antioxidant capacity (NADPH/NADP+). Pearson correlation coefficients for scatter plots were used to calculate r values with confidence intervals, and p values were determined by t-distribution.

### Increasing fl-E6 levels represses mitochondrial biogenesis and depletes antioxidant capacity

Fl-E6’s association with reduced mitochondrial mass and function prompted the question of whether manipulating E6 levels would alter mitochondrial biogenesis and oxidative phosphorylation. Because selective genetic silencing of E6 is hindered by E6 and E7 arising from the same transcript, we tested our hypothesis by increasing fl-E6 expression in VU147T and SCC154, the two HPV+ head and neck cancer lines with highest mitochondrial mass and lowest fl-E6 (Figure 3D). Lentiviral transduction of HPV16 fl-E6 was performed with a nonspliceable fl-E6 form (V42L) expressing a hemagglutinin (HA) tag (25) and used to generate three fl-E6-overexpressing clones and three vector control clones for each cell line. The three clones expressed fl-E6 mRNA at 60–300-fold (Supplemental Figure 3A) over endogenous levels, thus modeling the range of differences between the human tumors in TCGA with highest vs. lowest fl-E6 (Figure 3A, right). Fl-E6 overexpression did not alter *in vitro* growth (Supplemental Figure 3B) but produced the anticipated decrease in levels of E6’s canonical p53 target (Figure 4A). Increasing fl-E6 broadly repressed mRNA levels for 6 genes involved in oxidative metabolism: *SDHB* (Complex II), *UQCRC2* (Complex III), *COX II* (Complex IV), *ATP5A* (Complex V), *ACADM1* (acyl-CoA dehydrogenase medium chain-1), and *MDH1* (Malate dehydrogenase-1) (Supplemental Figure 3C). Similarly, five mitochondrial components examined by western blot using a commercial antibody cocktail *(NDUFB8, SDHB, UQCRC2, COX II* and *ATP5A)* were reduced by fl-E6 overexpression (Figure 4A, Supplemental Figure 3D). To determine whether these effects were generalizable beyond malignancy, fl-E6 was transduced by lentivirus into N-tert/1 keratinocytes expressing E7 (N-tert/E7 keratinocytes) (26). Three biological replicates expressing fl-E6 did not show altered *in vitro* growth (Supplemental Figures 3F and 3G), but reduction in expression of mitochondrial components was observed at the mRNA and protein level (Supplemental Figures 3H and 3I). Further characterization of VU147T and SCC154 cells overexpressing fl-E6 confirmed reductions in *MTCO1/B2M* (Figure 4B) and basal OCR (Figure 4C). Increased fl-E6 levels also depleted mitochondrial antioxidant capacity, as reflected by reduced NADPH/NADP+ ratio (Figure 4D). These results demonstrate fl-E6’s ability to attenuate mitochondrial mass, oxidative phosphorylation, and antioxidant capacity in HPV+ cancer cells and non-transformed cells expressing E7.

**Figure 4.**
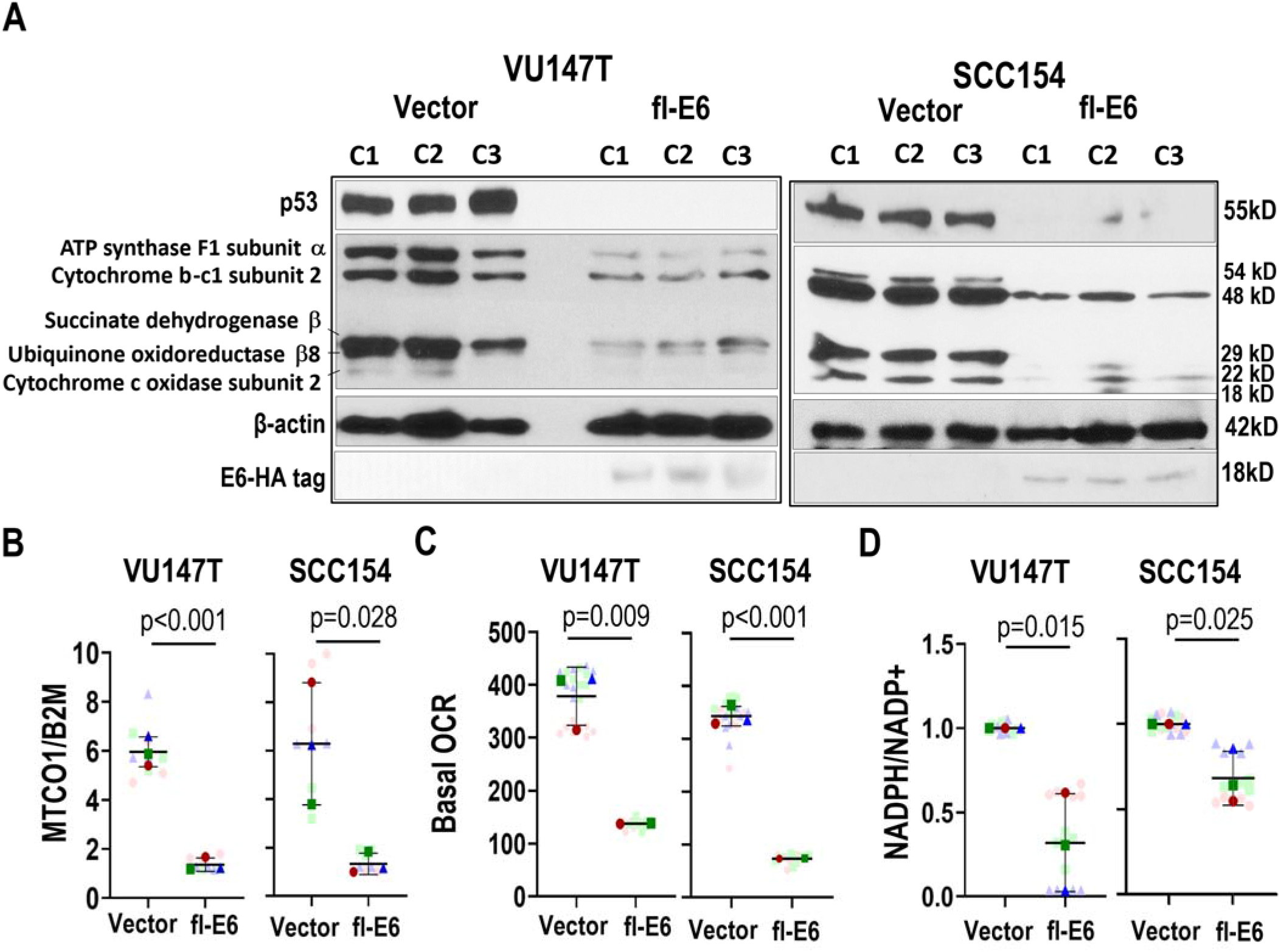
Increasing fl-E6 levels represses mitochondrial biogenesis and antioxidant capacity. **(A)** Western blot of 3 lentiviral fl-E6 transduced clones and 3 vector control clones showing levels of HA-tagged E6, p53, and five electron transport chain components. The effect of lentiviral fl-E6 expression is shown for **(B)** mitochondrial mass (*MTCO1/B2M* ratio by DNA qPCR), **(C)** basal OCR (Seahorse assay), and **(D)** antioxidant capacity (NADPH/NADP+ by enzyme cycling-based colorimetric assay). Super-plots display mean ± SEM of 3 technical replicates for each of three biologic replicates. Different colors represent biologic replicates. Light colors represent technical replicate values and corresponding dark colors represent their means. P values are based on unpaired t tests.

### Depletion of mitochondrial antioxidant capacity by fl-E6 potentiates therapy responses

The efficacy of radiation and cisplatin for many HPV+ OPSCCs indicates their limited capacity to maintain redox balance upon increasing ROS. We hypothesized that depletion of mitochondrial antioxidant capacity by fl-E6 contributes to therapy sensitivity in this setting. To test this possibility, radiation dose responses were measured in colony forming assays for VU147T and SCC154 cells, which contain low endogenous fl-E6. Lentiviral fl-E6 overexpression enhanced radiation dose responses (Supplemental Figure 4A) and decreased LD_50_ (Figure 5A). The same effect occurred with a chemical ROS inducer, 2,3-dimethoxy-1,4-naphthoquinone (DMNQ) (Figure 5B, Supplemental Figure 4B). Comparable effects of fl-E6 on cisplatin dose responses (Supplemental Figure 4B) and IC_50_ were shown by WST-1 assay (Figure 5C) and confirmed using a BrdU incorporation assay (Supplemental Figure 4C), which avoids potential for mitochondrial mass differences to confound cell number measurement in WST-1 assays. Because cisplatin may act by multiple mechanisms beyond ROS induction (27), we tested whether cisplatin sensitization by fl-E6 depended on mitochondrial antioxidant capacity depletion. Mito-TEMPO, a reagent that increases antioxidant capacity selectively in mitochondria at a 10μM dose (28), reversed the cisplatin sensitivity created in VU147T and SCC154 cells by fl-E6 (Figure 5D, Supplemental Figure S4D). Effects of increased fl-E6 on *in-vivo* growth and cisplatin responses were evaluated in xenografts of SCC154 and VU147T*. In-vivo* growth was maintained by both cell lines after lentiviral fl-E6 transduction but, in contrast to *in vitro* (Supplemental Figure 3B), slowed to variable extents (Figure 5E). Solid tumors from fl-E6-transduced cells showed fl-E6 mRNA expression at 15–60-fold the endogenous levels and maintained repression of oxidative metabolic transcript expression (Supplemental Figure 4E). Control tumors continued growth during treatment, confirming cisplatin resistance of these two low fl-E6 cell lines *in vivo*, and stable Fl-E6 expression markedly sensitized both to cisplatin treatment (Figure 5F, Supplemental Figure 4F). These results provide evidence that antioxidant capacity depletion by high fl-E6 levels contributes to the treatment sensitivity in HPV+ cancer; they further suggest that attenuated fl-E6 expression confers therapy resistance to certain tumors and enhances *in vivo* growth by mitigating oxidative stress.

**Figure 5.**
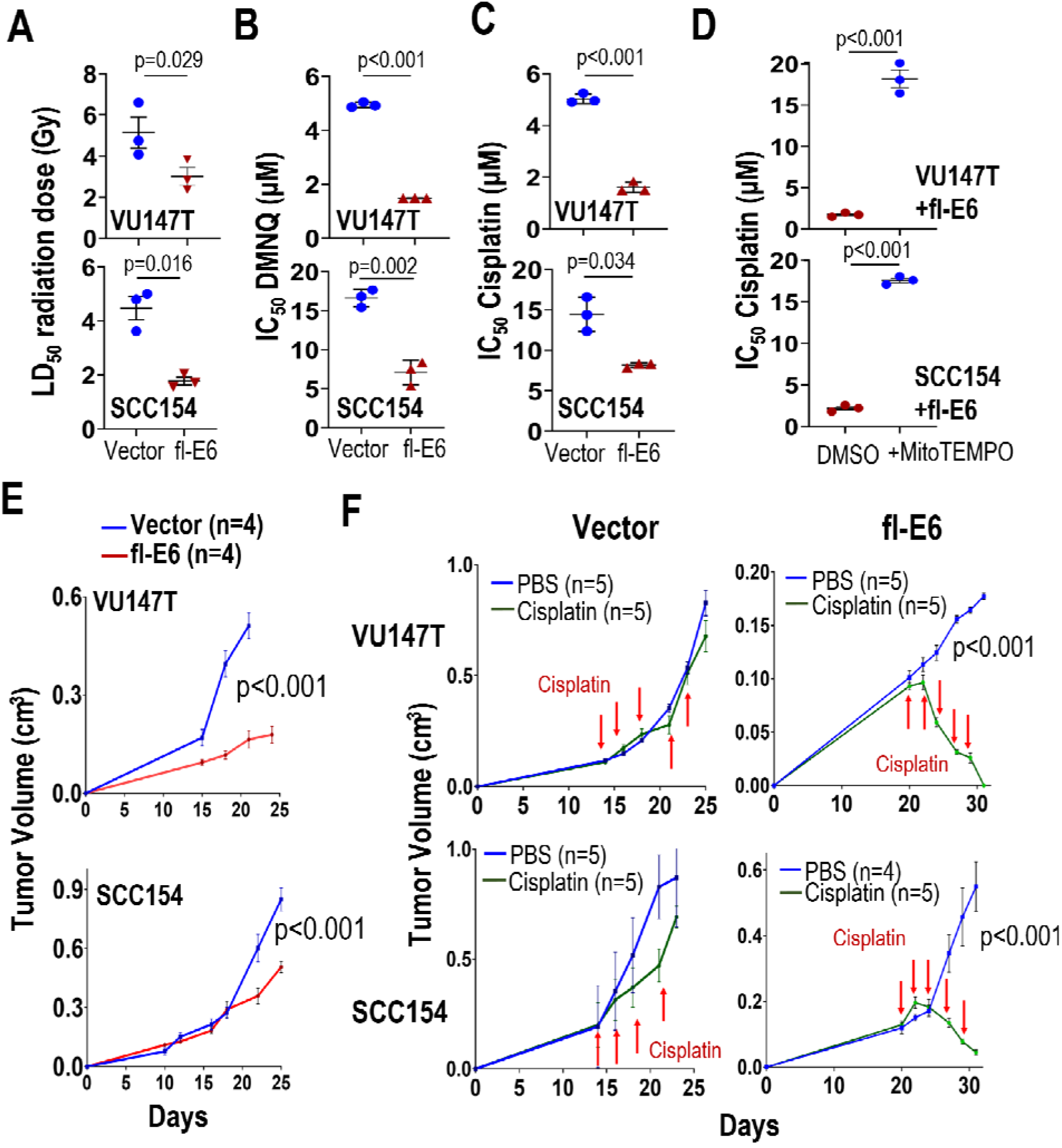
Depletion of mitochondrial antioxidant capacity by fl-E6 potentiates therapy responses. *In vitro* LD_50_ and IC_50_ were measured by simple logistic regression. **(A)** LD_50_ for radiation response is derived from number of cells surviving 10 days after 2, 4, and 6Gy. IC_50_ is quantified by WST-1 assay upon treatment with **(B)** DMNQ, **(C)** cisplatin, and by BrDu assay upon treatment with **(D)** cisplatin +/−10μM MitoTEMPO. Values represent mean ± SEM for 3 biological replicates. P values are from unpaired t tests. **(E)** *In vivo* subcutaneous tumor growth is shown in presence and absence of lentiviral fl-E6 expression. **(F)** treatment responses for both conditions are shown during intraperitoneal treatment with 1 mg/kg cisplatin 3x/wk for 2wks. P values of interaction were based on two-way ANOVA.

### Fl-E6 represses the PGC-1α/ERRα axis by reducing p53-dependent PGC-1α promoter activation

The central role of transcriptional coactivator PGC-1α and its homologue PGC-1β in mitochondrial biogenesis led to our testing whether fl-E6 negatively regulates their expression. Increasing fl-E6 expression in VU147T and SCC154 cells reduced mRNA levels for PGC-1α (Figure 6A) but not PGC-1β (Supplemental Figure 5A) and concomitantly decreased PGC-1α protein (Figure 6B, Supplemental Figure 5B). Levels of ERRα, PGC-1α’s obligate binding partner whose expression is positively regulated by PGC-1α in a feed-forward loop (16), also decreased (Supplemental Figure 5C). PGC-1α levels similarly decreased in the N-tert/E7 keratinocytes with fl-E6 expression (Supplemental Figures 5D and 5E), showing E6’s effect on PGC-1α to be generalizable to other cell contexts. Therefore, effects of fl-E6 on PGC-1α gene transcription were assessed using a luciferase reporter linked to the PGC-1α promoter. The reporter indicated a 50–75% decrease in PGC-1α promoter activity in VU147T and SCC154 cells upon fl-E6 overexpression (Figure 6C), consistent with p53’s direct positive regulation of the PGC-1α promoter in cancer cells (15). To discriminate among the multiple targets of E6 that might impact PGC-1α transcription, the same reporter was transfected into HEK293 cells along with mutant forms of E6 (Figure 6D). As anticipated, the non-spliceable fl-E6 construct and HPV16 wild type E6 comparably repressed reporter function, whereas the most common spliced form (E6*I) did not. PGC-1α promoter repression was maintained upon deleting the C-terminal PDZ-binding motif (E6Δ146–151) (29), showing E6’s effect to be independent of the multiple E6 target interactions mediated by PDZ domains. By contrast, E6 mutants with reduced capacity to bind p53 (8S9A10T) or reduced E6AP interaction (I128T) had reduced ability to repress the reporter. Because 8S9A10T and I128T only partly prevent p53 degradation and may not be p53-selective among E6’s targets (30), p53’s role was further evaluated using congenic HCT116 cells that are p53 wild type vs. null. The p53^−/−^ background reduced reporter activity relative to that in p53^+/+^ HCT116 cells, and fl-E6 repressed reporter function in p53^+/+^ cells but not in p53^−/−^ cells (Figure 6E). These findings demonstrate fl-E6’s capacity to repress PGC-1α expression and indicate a large component of this activity to be mediated by reduced p53.

**Figure 6.**
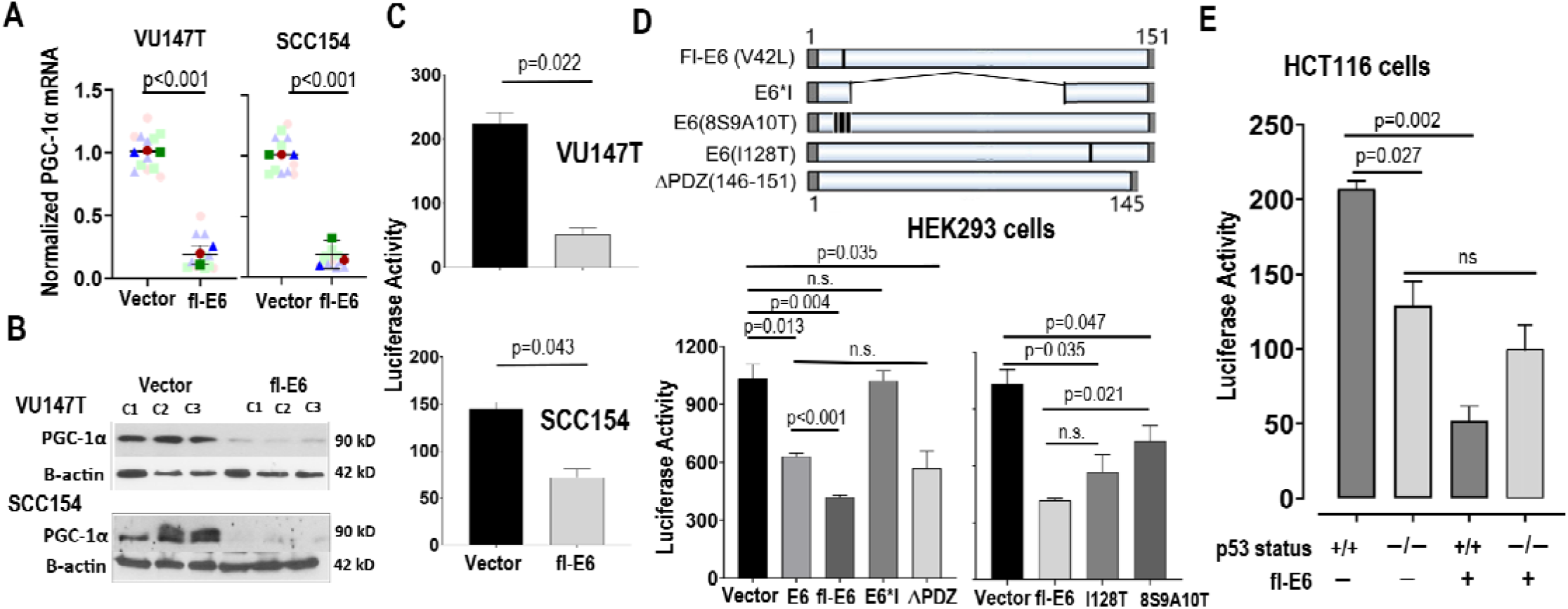
Fl-E6 represses PGC-1α levels by reducing p53-dependent PGC-1α promoter activity. **(A)** RT-qPCR for PGC-1α normalized to 18S after lentiviral fl-E6 expression in HPV+ cancer cell lines**. S**uperplots represent mean ± SEM for 3 biologic replicates with 4 technical replicates. P values are from unpaired t tests. **(B)** Western blot for PGC-1α in 3 lentiviral fl-E6 transfected clones and 3 vector control clones. **(C)** PGC-1α promoter-driven luciferase reporter activity in HPV+ cancer cell lines in presence and absence of lentiviral fl-E6 expression. **(D)** HEK293 cells co-transfected with the PGC-1α promoter-driven luciferase reporter plus the shown mutants of HPV16 E6 (top) before measuring luciferase activity (bottom). **(E)** *TP53* WT and null HCT116 cells were co-transfected with the PGC-1α promoter-driven luciferase reporter plus fl-E6 or empty vector. Reporter activity is normalized using background from a vector control reporter and expressed as mean±S.E. for 3 technical replicates. All results are representative of at least three independent experiments. P values were based on paired t-test.

### PGC-1α/ERRα axis activation in HPV+ OPSCCs with high mitochondrial mass and cisplatin resistance

Our observation that increasing E6 represses mitochondrial biogenesis by reducing p53-dependent PGC-1α expression led to us to evaluate this phenomenon for physiologic relevance in PDXs and patients, where fl-E6 levels vary widely. PGC-1α mRNA in the patient cohorts and PDX panel was captured by RNAseq at very low levels, often not meeting the detection threshold, and thus was instead quantified in the PDXs by qRT-PCR. As predicted, PGC-1α mRNA showed positive linear correlation with the number of upregulated Hallmark_Oxidative_Phosphorylation transcripts and *MTCO1/B2M* ratio (Figure 7A). Likewise, PGC-1α mRNA in the PDXs correlated negatively with fl-E6 mRNA (Figure 7B, left) and positively with protein levels of E6’s degradation target p53 (Figure 7B, middle, Supplemental Figure 5F). The predicted negative correlation between PGC-1α expression and cisplatin resistance measured by rate-based T/C was confirmed in the PDXs (Figure 7B, right). To overcome technical inability to detect PGC-1α in the clinical cohorts by RNAseq, we examined as a surrogate marker its obligate binding partner ERRα, whose role as transcriptional target of PGC-1α/ERRα creates a feed-forward loop that amplifies PGC-1α’s activity (16). ERRα was quantifiable in the cohorts in Figure 1 where RNAseq was performed using frozen tissue (TCGA and JHU) but not FFPE (VU), thus providing a readout for PGC-1α/ERRα axis activation. A positive linear correlation was observed between ERRα levels and number of up-regulated Hallmark_Oxidative_Phosphorylation transcripts in the PDX panel (Figure 7C, left) and both patient cohorts (Figure 7C, right). Moreover, ERRα mRNA correlated negatively with fl-E6 in TCGA, the only cohort with fl-E6 annotation (Figure 7D). Lastly, ERRα mRNA levels positively correlated with the number of upregulated transcripts in the Hallmark_P53_Pathway gene set in both cohorts (Figure 7E), with upregulation defined by the same criteria used for Hallmark_Oxidative_Phosphorylation genes in Figure 1. In context of our mechanistic findings in cell lines, these concordant observations in PDXs and patients support the concept that variable fl-E6 levels among HPV+ OPSCCs lead to differential PGC-1α/ERRα pathway activity by a p53-mediated mechanism, in turn causing variability in mitochondrial mass that affects therapy responses.

**Figure 7.**
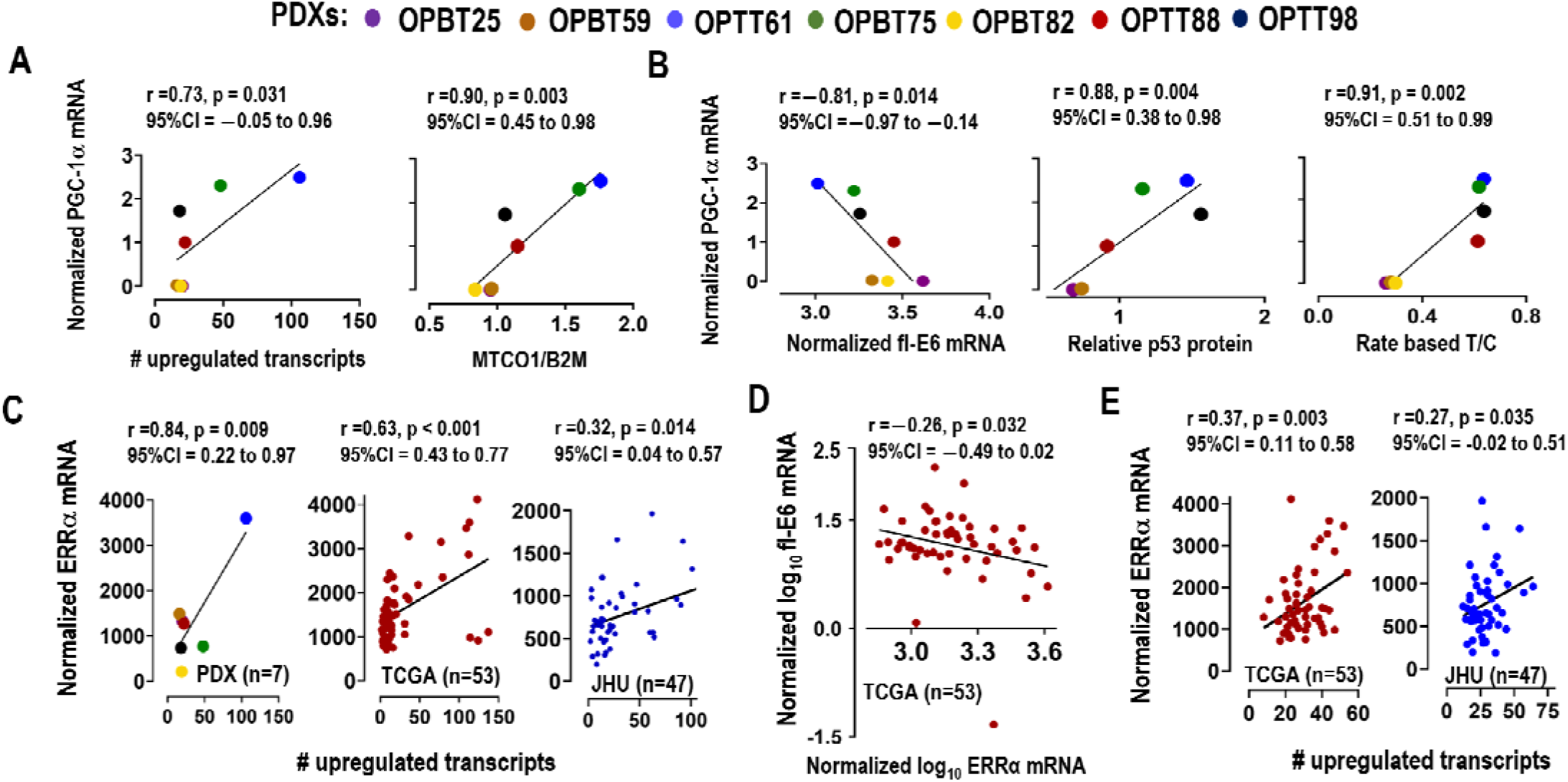
PGC-1α/ERRα axis activation in tumors with high mitochondrial mass and therapy resistance. PGC-1α was measured in HPV+ OPSCC PDXs by RT-qPCR normalized to 18S. All other transcripts were measured by RNAseq. Pearson correlation coefficients for scatter plots were used to calculate r values with confidence intervals, and p values were determined by t-distribution. **(A)** PDX PGC-1α mRNA vs. the number of upregulated Hallmark_Oxidative_Phosphorylation transcripts (left) and vs. mitochondrial mass (right) *(MTCO1/B2M* by DNA qPCR). **(B)** PDX PGC-1α vs. fl-E6 mRNA (left), p53 levels from western blot densitometry (middle), and *in vivo* cisplatin response by rate-based T/C (right). **(C)** ERRα mRNA vs. number of up-regulated Hallmark_Oxidative_Phosphorylation transcripts in the PDXs (left) and the HPV+ OPSCCs in TCGA cohort (middle) and the JHU cohort (right). **(D)** fl-E6 vs. ERRα mRNA in TCGA cohort. **(E)** ERRα mRNA vs. number of up-regulated Hallmark_P53_Pathway transcripts in TCGA cohort (left) and the JHU cohort (right).

## DISCUSSION

The impact of variation in viral oncoprotein levels on clinical behavior of HPV+ cancers has been unclear to date. Furthermore, unknown molecular mechanisms distinguishing favorable vs. unfavorable responses to therapy have hindered personalizing management for HPV+ OPSCCs. Our results reveal a novel ability of E6-mediated p53 downregulation to deplete mitochondrial antioxidant capacity and thus sensitize HPV+ OPSCCs to standard therapy, which relies upon induction of oxidative stress. We provide evidence that differing fl-E6 levels across HPV+ cancers variably repress the PGC-1α/ERRα pathway, leading to diversity in mitochondrial mass that likely affects treatment response. This finding provides a novel mechanism by which levels of a viral oncoprotein may govern treatment outcomes in HPV+ OPSCC. Therapy resistance may prove prospectively identifiable by measuring fl-E6 levels and downstream components of the PGC-1α/ERRα axis and be mitigated through therapeutically targetable nodes in that pathway.

Our findings add to understanding of how variable HPV oncogene expression effects cancer clinical behavior. HPV E2 disruption by viral genome integration in a subset of cancers is traditionally thought to derepress E6/E7 expression, and there is some evidence suggesting E2 disruption portends worse prognosis (31). Increased E1∧E4 mRNA (32) and decreased E2F target gene dysregulation downstream of E7 (22) have also been linked to worse survival in small clinical cohorts, albeit without mechanistic basis. Increased E6 splicing is observed in cancer relative to normal viral replication (33) and may indicate that the fl-E6 levels best serving malignant progression are lower those that serve the viral life cycle. Similarly, chronic *in vitro* cisplatin exposure is reported to reduce fl-E6 expression (34) and may represent adaptation to mitigate oxidative stress. Taken together with our findings, these prior observations suggest that certain HPV+ cancers achieve competitive advantage through partially preserving p53 function by down-regulating fl-E6.

The p53-mediated transcriptional upregulation of PGC-1α observed in other cancer types (15) agrees with our observed mechanism of E6 repressing mitochondrial biogenesis but may not fully account for E6’s effects in this context. Our observations notably diverge from findings in mouse embryonic fibroblasts, where p53 was activated by telomere shortening and directly bound and repressed PGC-1α, thus contributing to growth arrest, senescence, and apoptosis (35). By contrast, both wild type and mutant p53 can act as positive regulatory cofactors for PGC-1α at promoters of its target genes in multiple cancer types (36, 37). Increased p53 also drives expression of sestrins (38), which mediate PGC-1α activation by stimulating its phosphorylation by AMPK (39). In addition, p53 supports mitochondrial DNA replication via interaction with mitochondrial transcription factor A (40). P53-independent mechanisms by which E6 might suppress mitochondrial biogenesis also exist and include E6’s degradation of p300/CBP (41), whose coactivation of CREB contributes to PGC-1α promoter activity (42). E6 also targets TIP60, another acetyltransferase whose substrate BRD4 activates a super-enhancer of PGC-1α expression (43). These mechanisms highlight AMPK, CREB, p300/CBP, TIP60, and BRD4 as targetable nodes upstream of PGC-1α/ERRα with potential utility for phenocopying E6’s desirable effects on therapy response.

The PGC-1α/ERRα complex and its myriad downstream effects on mitochondrial metabolism offer additional potential targets for treatment-resistant HPV+ OPSCCs. Pharmacologic ERRα inhibition was recently achieved using diaryl ether-based thiazolidinediones, which interfere with its DNA binding (44). Targeting p300/CBP’s co-activator function for PGC-1α (45) offers another approach to PGC-1α/ERRα axis repression and may bring an additional benefit of repressing DNA repair in certain HPV+ tumors with p300-activating mutations (46). Furthermore, inhibitors of electron transport chain complexes, TCA cycle enzymes, and glutamine metabolism in clinical trials for various treatment-refractory cancers (47) may have value for a subset of HPV+ OPSCCs of similar metabolic phenotype. Although aggressively targeting oxidative metabolism in the clinic remains hampered by toxicity, the severely impaired DNA damage repair phenotype in HPV+ OPSCCs (3) may widen the therapeutic window by allowing responses at lower drug doses than those needed for other cancers.

E6 levels and mitochondrial mass are related to other features of HPV+ OPSCCs being explored as biomarkers. Image-based quantification of hypoxia as a source of radio-resistance is in clinical trials to guide therapy de-escalation for HPV+ OPSCC (48). Because hypoxia induces oxidative stress, tumors with the antioxidant capacity to survive hypoxia may exploit the same antioxidant mechanisms to resist treatment. The benefit of increased mitochondrial function in this context is consistent with decreases in E6/E7 expression and p53 degradation (49) observed under hypoxia in HPV+ cervical cancer cell lines. There is also evidence that genetic silencing of negative regulators of NRF2, a driver of antioxidant gene transcription with downstream and upstream relationships to PGC-1α, confers poor prognosis in HPV+ cancers (12). Therefore, fl-E6 levels may merit joint consideration with NRF2 regulation, hypoxia, and other features linked to antioxidant capacity in developing multi-marker predictors of therapy response.

## METHODS

### Patient cohorts

Data for the Head and Neck Squamous Cell Carcinoma TCGA cohort were downloaded from the Genomic Data Commons (project TCGA-HNSCC) and Broad GDAC Firebrowse repository. 53 HPV+ cases were selected from this broader TCGA cohort based on oropharyngeal primary site and previous detection of high-risk HPV type E6/E7 transcript expression (17). RNAseq and clinical information for the 47 HPV+ OPSCCs in a Johns Hopkins University (JHU) cohort are previously described (20). RNAseq and clinical data were obtained for the 37 HPV+ OPSCCs treated with primary radiation plus chemotherapy within a published Vanderbilt University (VU) cohort (21).

### Bioinformatic analysis of TCGA RNAseq data

RNA-Seq reads within FastQ files for each TCGA OPSCC sample (n=53) were aligned to HPV genomes using Bowtie2 and samples with alignment selected and identified as containing either HPV 16, HPV 33, HPV 35, or HPV 56 genomes. Gene expression values were represented by the normalized reads of Reads Per Kilobase of transcript per Million mapped reads (RPKM). HPV genome sequences and annotations with gene coordinates were obtained from RefSeq (https://www.ncbi.nlm.nih.gov/refseq/). Reads with mapped coordinates corresponding to regions specific for each gene were counted. Full-length E6 expression was quantitated in a region non-overlapping with E6*. The probe region specific to E6* overlaps full-length E6 and thus represents total E6. E6* is calculated as by subtracting full-length E6 to Total E6. HPV16 gene expression analysis used GenBank: K02718.1, with the following gene coordinates for quantitation: E6: 227–408, total E6: 83–226, E7: 562–858, E2: 2814–3331, E4: 3332–3619, and E5: 3863–4099. HPV33 gene expression analysis used GenBank: M12732.1, with the following gene coordinates for quantitation: E6: 232–413, total E6: 109–230, E7: 573–866, E2: 2814–3325, E4: 3326–3577, and E5: 3854–4081. HPV35 gene expression analysis used GenBank: X74477.1, with the following gene coordinates for quantitation: E6: 233–414, total E6: 110–231, E7: 562–861, E1: 868–2713, E2: 2882–3293, E4: 3294–3584 and E5: 3814–4065. HPV56 gene expression analysis used GenBank: X74483.1, with the following gene coordinates for quantitation: E6: 234–415, total E6: 102–232, E7: 572–889, E2: 2806–3221, E4: 3222–3577, and E5: 3854–4081.

### Growth and RNA sequencing of PDXs

The seven treatment-naïve HPV+ OPSCC PDXs used in this study were established and passaged as described by us in the subcutaneous flank of NSG mice (22). The PDXs were harvested for RNAseq in triplicate upon reaching 1cm^3^. Volumes were calculated as width^2^ × length/2. RNA isolation from 7 PDXs was performed with the Qiagen RNA Kit (Qiagen, Hilden, Germany). Sample quality was assessed using Agilent RNA 6000 Nano Reagent on Agilent 2100 Bioanalyzer (Agilent Technologies, Santa Clara, CA) and quantified by Qubit RNA HS assay (ThermoFisher, Waltham, MA). Ribosomal RNA depletion was performed with Ribozero rRNA Removal Kit, HMR (Illumina Inc., San Diego, CA). Library preparation was performed with the NEBNext Ultra II Non-directional Synthesis Module (New England Biolabs, Ipswich, MA). Samples were sequenced on Illumina NextSeq High Output sequencer and aligned to combined genome from hg38 (human), mm10 (mouse) and the high-risk HPV genomes (16, 18, 31, 33, 35, 39, 45, 51, 52, 56, 58, 59, 66 and 68) using STAR with default parameters. HPV sequences in all samples mapped to HPV16 exclusively. After STAR first-pass alignment, reads mapped to mm10 were removed, and reads aligned to human genome (hg38) and HPV genomes were retained. Raw counts for each viral transcript (E2, E4, E5, L2, L1, total-E6, E7 and E1∧E4) were obtained from reads mapped to each gene based on the alignment file. Fl-E6 was computed as the ratio of the average coverage level in the first intron (approximate full-length E6 level) divided by the average coverage level in the first exon (approximate all E6 transcript level). RnaSTAR was used to identify the first introns for E6*, which are, in the format of [donor-acceptor], [227–408] for HPV-16. The average coverage of read pairs within the intron (i.e., 227–408 for HPV-16) was divided by the average coverage of read pairs within exon1 (i.e., 82–227 for HPV-16), where coverage was computed by dividing the number of reads by the feature width. RNAseq data reported here was submitted to GEO record GSE193388.

### Cell lines and culture conditions

SCC090 (RRID:CVCL_1899), SCC152 (RRID:CVCL_0113), SCC154 (RRID:CVCL_2230), HEK293(RRID:CVCL_0045), HCT116 (RRID:CVCL_0291), and HCT116 p53-/-(RRID:CVCL_HD97) cells are from ATCC. UM-SCC047 (RRID:CVCL_7759) and UMSCC104 (RRID:CVCL_7712) are from Sigma-Aldrich.

UDSCC2 (RRID:CVCL_E325) and 93VU147T (RRID:CVCL_L895) cell lines were gifts from Silvio Gutkind (U. California, San Diego), and Hans Joenje (VU Medical Center, Netherlands), respectively. Cell lines were authenticated using the Identify Mapping Kit (Coriell, Camden, NJ). They were cultured in DMEM-F12 (Gibco, Gaithersburg, MD), 10% heat-inactivated FBS, 1% Pen-Strep, and 1% nonessential amino acids. N/Tert-1/E7 cells were cultured in keratinocyte serum-free medium (Thermo-Fisher, Waltham, MA).

### Quantitative real-time PCR and western blotting

Total RNA was extracted using the RNeasy Plus Micro Kit (Qiagen, Hilden, Germany). DNase I (AMPD1, Sigma-Aldrich, St. Louis, MO) was used to remove genomic DNA from RNA samples. Reverse transcription was performed with oligo-dT plus random decamer primers using Superscript II (Thermo-Fisher). Quantitative PCR was performed with SYBR green master mix (Thermo-Fisher) using Step-one Plus Real-Time PCR. Primers are in Supplemental Table 2. Western blots were performed with luminol-based chemiluminescent substrate (Bio-Rad, Hercules, CA), and bands in X-ray films were quantified with ImageJ software (RRID:SCR_003070). Antibodies are detailed in Supplemental Table 3.

### Lentiviral transduction

The pLentiN-16E6no* (RRID:Addgene_37445) plasmid containing full-length E6 and its pLentiN (RRID:Addgene_37444) vector backbone control were used to stably express E6. Lentiviral preparation and transduction were performed as described (25). Blasticidin selection at 2–5μg/ml was performed for 3–4 weeks before subculturing and plating at limiting dilution in 96 well plates. N/Tert-E7 cells were infected with the same control and E6 viruses before blasticidin selection and propagation.

### Luciferase assays

HPV16 E6 mutant constructs 16E6, 16E6(V42L), 16E6–8S9A10T, 16E6-I128T, and 16E6* are previously described (29). The pGL3 luciferase reporter plasmid was used as the backbone for 2423bp human PGC-1α promoter. Plasmids were co-transfected using Lipofectamine 2000 (Thermo-Fisher). Fold changes were determined by measuring the ratio of firefly to Renilla luciferase activity and comparing it to the ratio for control vector-transfected cells. Plasmid details are in Supplemental Table 4.

### Evaluating mitochondrial mass and function

Mitochondrial DNA content was determined by qPCR as described (23). Oxygen consumption rate (OCR) was measured on the Seahorse Bioscience™ XFe96 Extracellular Flux Analyzer (Seahorse Bioscience, North Billerica, MA) per manufacturer protocol using 0.001mg/ml oligomycin, 2.5μM carbonyl cyanide p-trifluoro-methoxyphenyl hydrazone, and 2μM each of antimycin and rotenone (Sigma-Aldrich). Total protein per well was determined after with the Pierce™ BCA Protein Assay Kit (Thermo-Fisher). Results were analyzed using XFe Wave software (Seahorse Bioscience). 1×10^5^ cells were used to measure NADPH and NADP^+^ levels using a NADP+/NADPH colorimetric quantitation kit (Sigma-Aldrich).

### Pharmacologic and radiation therapy experiments

Cell viability after 72 hours treatment with cisplatin, DMNQ, and/or MitoTEMPO (Sigma-Aldrich) was determined by WST-1 and/or BrDu assays (Sigma-Aldrich). Cisplatin dissolved in saline was administered intraperitoneally *in vivo* at 2mg/kg. For irradiation, cells were harvested during exponential growth and plated at 2000 cells/ml in 60mm dishes containing 2ml media. After 12 hours, they received 2, 4, or 6 Gy using a Gamma cell 40 irradiator. Media was changed post-irradiation. Colonies were stained with crystal violet and counted 10 days later.

### Statistics

When comparing two groups for cell viability or gene expression, significance was calculated using two-tailed unpaired t-tests. Pearson correlation coefficients were based on analyzing the means. Two-way ANOVA was used when comparing xenograft growth curves. Log-rank tests were used for Kaplan-Meier analyses. Tests used are indicated in figure legends. Analyses were performed using Prism (GraphPad Software, San Diego, CA).

### Study approval

Mouse experiments were performed under Wistar Institute IACUC protocols #201166 and #201178. PDXs were previously established under University of Pennsylvania IRB-approved protocol #417200 “Head and Neck Cancer Specimen Bank” (PI: D. Basu) by signing a combined informed consent and HIPAA form for use of tissue for research. All clinical data shown is obtained from de-identified, publicly available datasets.

## AUTHOR CONTRIBUTIONS

M.S. and D.B. conceived the study. D.B., M.S., E.W., I.M., and B.W. wrote and edited the manuscript and/or designed figures and tables. M.S., P.R., L.L., and V.S. performed experiments. P.G., P.R., M.S., X.W., X.L, and B.W. performed bioinformatic and/or statistical analyses. D.B., E.W., P.G., D.K., and H.N. supervised the study and data analysis. D.B, J.J., R.B., X.W., and X.L. analyzed clinical and/or pathologic data. All authors read and approved the final manuscript.

## ACKNOWLEDGEMENTS

This study was supported by NIH R01-DE027185 (DB, PG), P30-DK050306 (DB), R01-HL058493 (DK, LL), R01-DE026471 (XL, XW), P01-CA098101 (HN), R01-AA026297 (HN), P30-CA013696 (HN) and R03-DE026230 (BW, IM). We thank Dr. Roger Cohen for pre-submission review of the manuscript.

## Notes

**Conflict of interest statement:** The authors have declared that no conflict of interest exists.

### Competing Interest Statement

The authors have declared no competing interest.

