## Supplemental data file for "Human papilloma virus E6 regulates therapy responses in oropharyngeal cancer by repressing the PGC-1α/ERRα axis"

**INDEX**

**Supplemental Figure 1.** Characterization of mitochondrial function and cisplatin response in HPV+ PDXs and cell lines

**Supplemental Figure 2.** Lack of associations between mitochondrial mass and levels of other HPV oncogenic transcripts in TCGA and the PDX panel

**Supplemental Figure 3.** Metabolic effects of increasing fl-E6 mRNA expression on VU147T cells, SCC154 cells, and N-tert/E7 keratinocytes

**Supplemental Figure 4.** Effect of increased fl-E6 on treatment sensitization of VU147T and SCC154 cells

**Supplemental Figure 5.** Effects of increased fl-E6 expression on the PGC-1α/ERRα axis in cancer cell lines, N-tert/E7 keratinocytes, and HEK293 cells

**Supplemental Table 1.** Patient characteristics in TCGA, JHU, and VU cohorts

**Supplemental Table 2.** Primers used for qPCR analysis

**Supplemental Table 3.** Antibodies used in the study

**Supplemental Table 4.** Plasmids used in the study

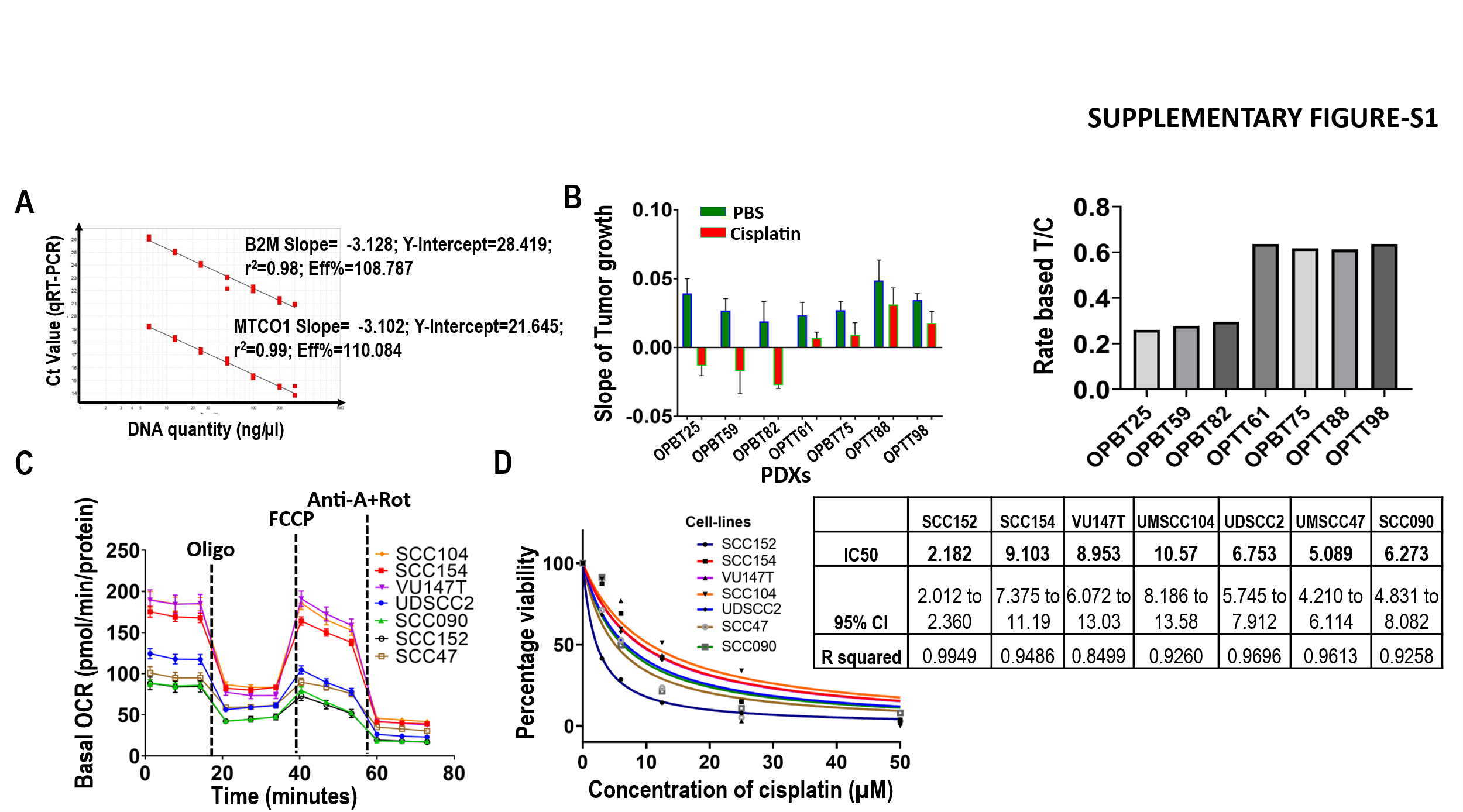

**Supplemental Figure 1.** **Characterization of mitochondrial function and cisplatin response in HPV+ PDXs and cell lines. (A)** Standard curves used to determine *MTCO1/B2M* ratio by DNA qPCR. **(B)** Tumor growth rate slopes used to calculate rate-based T/C value [25] for PDXs after intraperitoneal administration of PBS or Cisplatin 3 times/week for 2 weeks. At least 4 biological replicates were used for each condition, and tumor volumes were measured at times of cisplatin administration. **(C)** Seahorse Assay for cell line panel showing normalized OCR profile after addition of oligomycin (Oligo, 0.001 mg/ml), carbonyl cyanide p-trifluoro-methoxyphenyl hydrazone (FCCP, 2.5 µM), and antimycin + rotenone (Anti-A+Rot, 2 µM each). **(D)** Dose response curve for cisplatin vs. normalized cell viability by WST assay after 48 hours treatment. IC50 with 95% confidence intervals (CI) and R^2^ were obtained from simple logarithmic regression. Data points represent mean ± SEM.

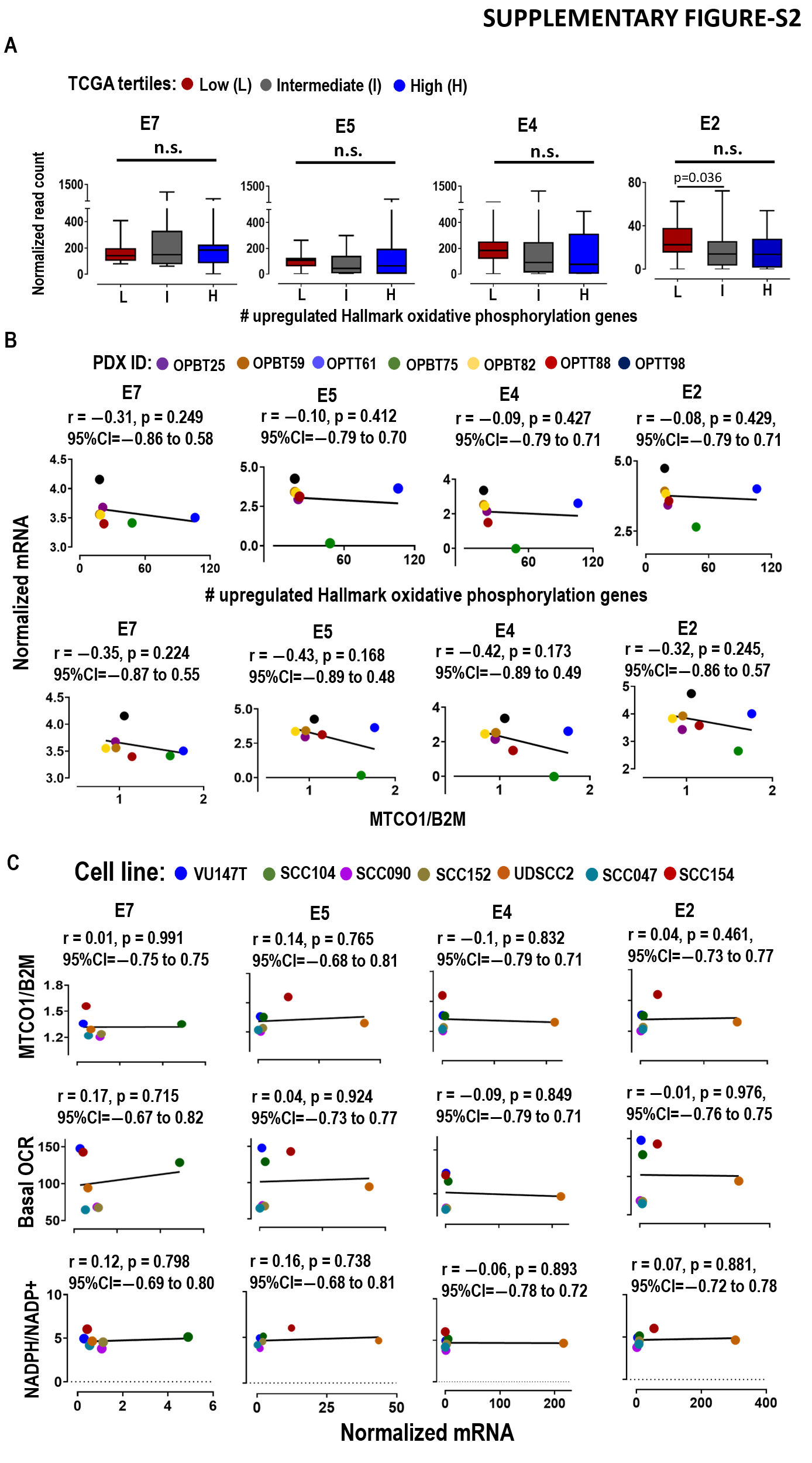

**Supplemental Figure 2. Lack of associations between mitochondrial mass and levels of other HPV oncogenic transcripts in TCGA and the PDX panel. (A)** RNAseq evaluating E7, E5, E4, and E2, for differential expression among tertiles of Hallmark Oxidative Phosphorylation gene expression in TCGA. p values calculated by Mann Whitney Test. **(B)** Scatter plots for the PDXs evaluating the same HPV transcript levels vs. number of up-regulated Hallmark Oxidative Phosphorylation transcripts (top) and mitochondrial mass (*MTCO1/B2M* by DNA qPCR) (bottom). **(C)** Scatter plots for HPV+ cell lines evaluating the same HPV transcript levels vs. mitochondrial mass (MTCO1/B2M DNA qPCR, above), basal OCR (middle) measured by Seahorse Assay and vs. NADPH/NADP+ (below) measured by enzyme cycling-based colorimetric assay. Pearson correlation coefficients were used to calculate r values with confidence intervals, p values determined by t-distribution.

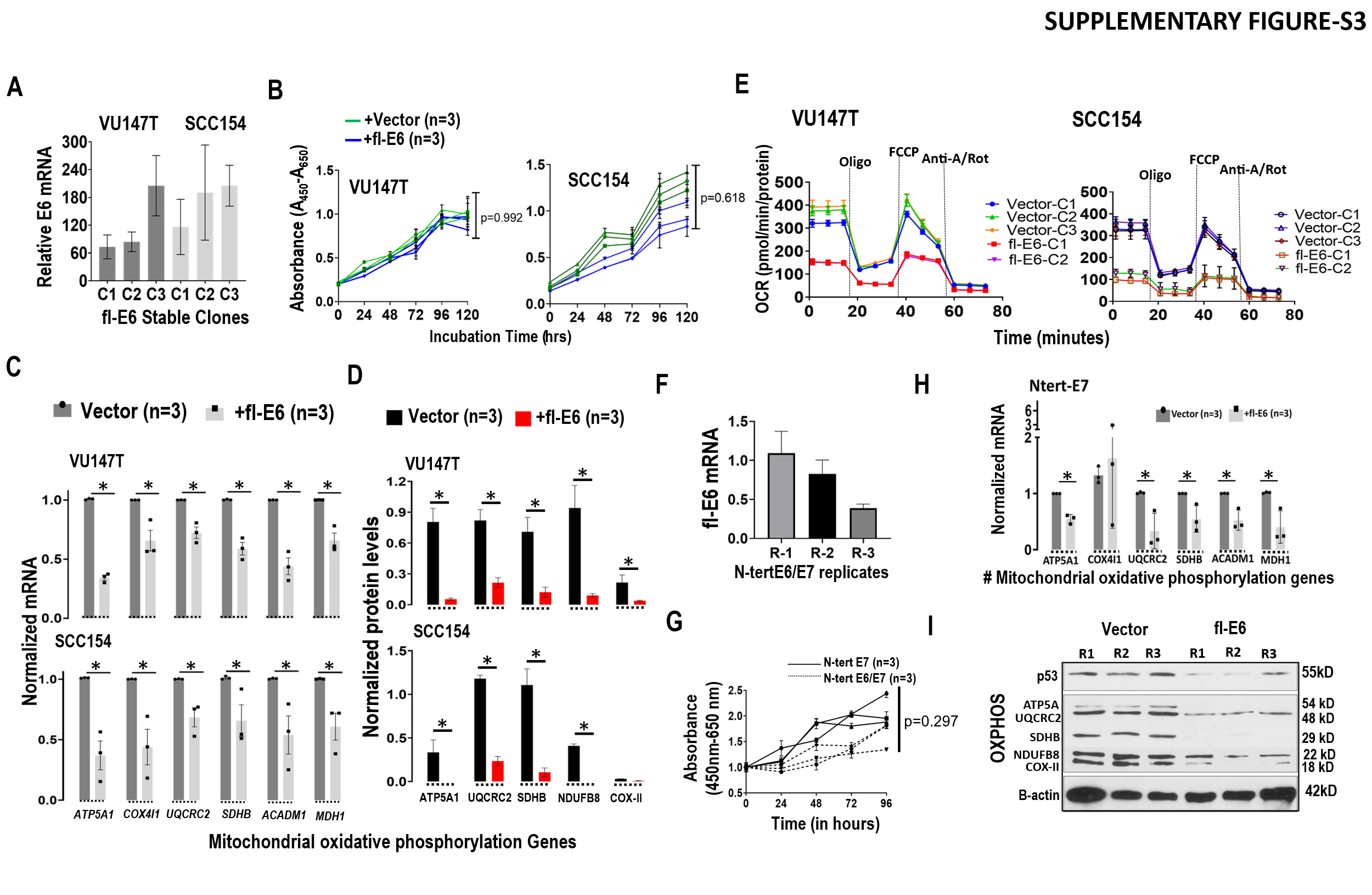

**Supplemental Figure 3. Metabolic effects of increasing fl-E6 mRNA expression on VU147T cells, SCC154 cells, and N-tert/E7 keratinocytes. (A)** RT-qPCR of E6 mRNA normalized to 18S in 3 lentiviral fl-E6 transfected clones and 3 vector control clones of SCC154 and VU147T. **(B)** Growth rate of SCC154 and VU147T upon stable fl-E6 expression by WST assay. p value is calculated using ANOVA. **(C)**RT-qPCR upon fl-E6 expression for 6 genes involved in oxidative phosphorylation, with 18S as internal control. **(D)** Densitometry of western blots for 5 mitochondrial electron transport chain proteins upon fl-E6 expression, normalized using β-actin. Bars represent mean ± SEM for three biologic replicates. p value calculated using unpaired t test. *p<0.05. **(E)** Seahorse Assay showing effect of fl-E6 expression on normalized OCR profile after addition of oligomycin (Oligo, 0.001 mg/ml), carbonyl cyanide p-trifluoro-methoxyphenyl hydrazone (FCCP, 2.5 µM), and antimycin + rotenone (Anti-A+Rot, 2 µM each). **(F)** RT-qPCR for fl-E6 mRNA normalized to 18S upon stable transfection of N-tert/E7 keratinocytes. **(G)** Effect of fl-E6 expression on growth rate of N-tert/E7 keratinocytes by WST assay, showing 3 biologic replicates per condition. p value by ANOVA. **h** RT-qPCR in N-tert/E7 keratinocytes +/- fl-E6 for 6 genes involved in oxidative phosphorylation normalized by 18S. Values represent mean ± SEM for three biologic replicates. p value by unpaired t test. *p<0.05. **i** Western blot of 3 biologic replicates of N-Tert/E7 cells+/-fl-E6 showing p53 and five electron transport chain components.

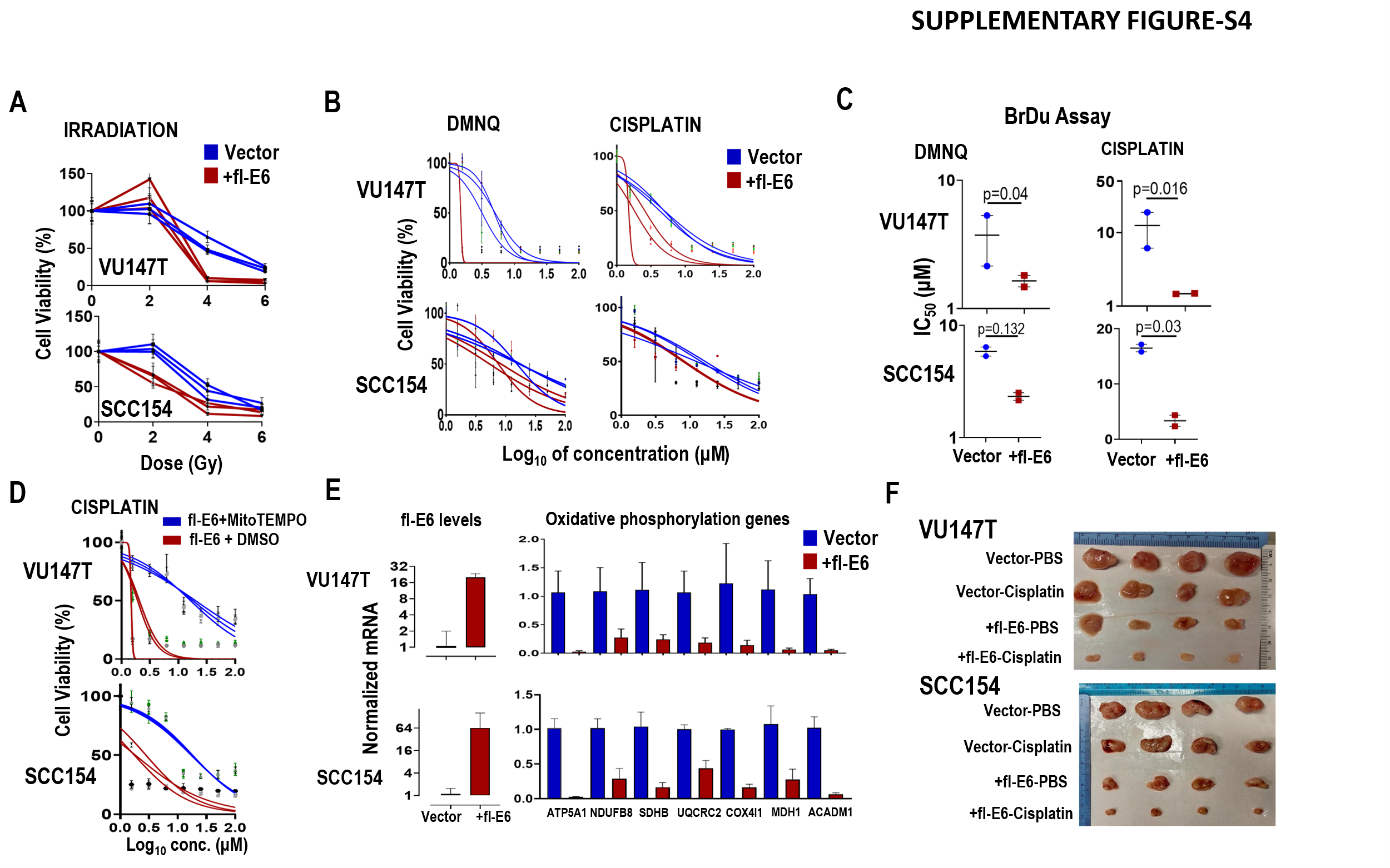

**Supplemental Figure 4. Effect of increased fl-E6 on treatment sensitization of VU147T and SCC154 cells.** *In vitro* IC_50_ were quantified by simple logistic regression. **(A)** % surviving cells 10 days after 2, 4, or 6 Gy radiation. **(B)** Dose responses to DMNQ (left) and cisplatin (right) based on % surviving cells after 48 hours of treatment using WST assay. **(C)** IC_50_ for DMNQ (left) and cisplatin (right) after 48 hours of treatment using BrDu colorimetric assay. **(D)** Dose responses to cisplatin +/- 10μM MitoTEMPO/DMSO after 48 hours of treatment using BrDu colorimetric assay. Each point on the survival curve and dose response represents the mean surviving fraction from at least three replicates. **(E)** VU147T and SCC154 cells +/- stable fl-E6 expression was grown subcutaneously *in vivo* for 2 weeks. qRT-PCR normalized to 18S shows expression of fl-E6 and 7 genes involved in oxidative phosphorylation in resulting solid tumors. Bars represent mean ± SEM for two solid tumors per group. **(F)** Image of solid tumors used to quantify *in vivo* responses to cisplatin in Figure 4F.

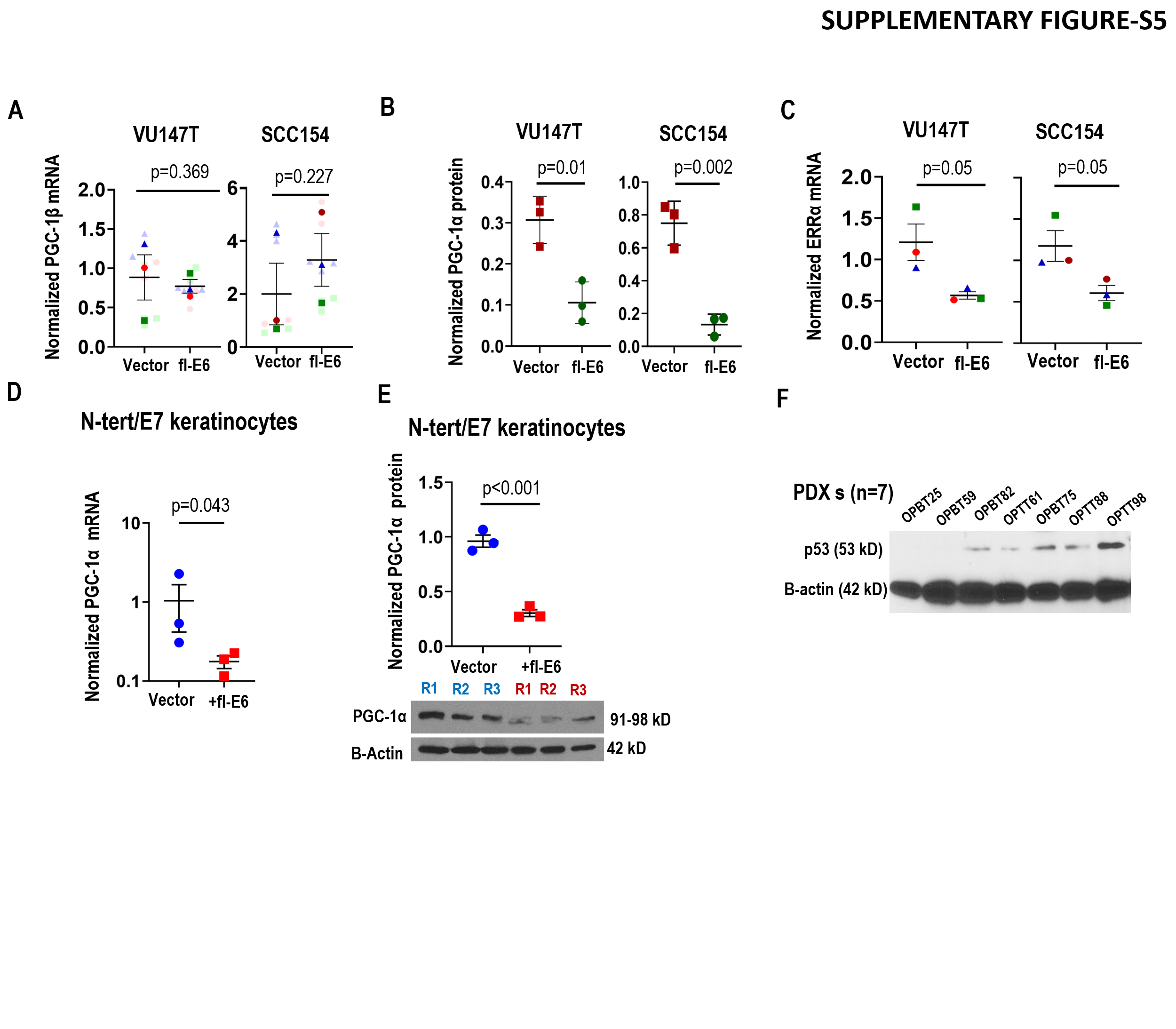

**Supplemental Figure 5. Effects of increased fl-E6 expression on the PGC-1α/ERRα axis in cancer cell lines, N-tert/E7 keratinocytes, and HEK293 cells.** **(A)** RT-qPCR of PGC-1β transcript normalized to 18S. **(B)** Western blot densitometry for PGC-1α normalized to β-actin. **(C)** RT-qPCR of ERRα transcript normalized to 18S in 3 lentiviral fl-E6 transfected clones and 3 vector control clones of SCC154 and VU147T. Plots represent mean ± SEM for 3 biologic replicates. p value calculated using unpaired t test. **(D)** RT-qPCR of PGC-1α transcript normalized to 18S. **(E)** Western blot of PGC-1α (left), with band densities (right) normalized to β-actin in 3 lentiviral fl-E6 transfected and 3 vector control replicates of N-tert/E7 keratinocytes. Plots represent mean ± SEM for the biologic replicates. p value calculated using unpaired t test. **(F)** Western blot of p53 and β-actin in PDX panel.

**Supplemental Table 1. Patient characteristics in TCGA, JHU, and VU cohorts**

| **Variable** | **Category** | **TCGA (n=53)** | **JHU**  **(n=47)** | **VU**  **(n=37)** |
| --- | --- | --- | --- | --- |
| **Median age (range)** |  | 56 (35-77) | 55 (35-75) | 58 (52-62) |
| **Gender, N (%)** | Male | 48 (90.6) | 42 (89.4) | 2 (5.4) |
|  | Female | 5 (9.4) | 5 (10.6) | 35 (94.6) |
| **Clinical T-stage†,**  **N (%)** | Early (Tx/T0–2) | 40 (75.5) | 35 (74.5) | 23 (62.2) |
|  | Advanced (T3–4) | 5 (9.4) | 12 (25.5) | 13 (35.1) |
|  | Unknown | 8 (15.1) | 0 (0) | 1 (2.7) |
| **Clinical N-stage†,**  **N (%)** | N0 | 7 (13.2) | 2 (4.2) | 3 (8.1) |
|  | N1 | 5 (9.4) | 5 (10.6) | 6 (16.2) |
|  | N2a | 2 (3.8) | 10 (21.3) | 5 (13.5) |
|  | N2b | 9 (16.9) | 20 (42.6) | 14 (37.8) |
|  | N2c | 0 (0) | 2 (4.2) | 8 (21.6) |
|  | N3 | 1 (2) | 2 (4.2) | 0 (0) |
|  | Unknown | 29 (54.7) | 6 (12.8) | 0 (0) |
| **Clinical M-stage^†^,**  **N (%)** | M0 | 14 (26.4) | 47 (100) | 35 (94.6) |
|  | M1 | 0 (0) | 0 (0) | 0 (0) |
|  | Unknown | 39 (73.6) | 0 (0) | 2 (5.4) |
| **Clinical overall stage^†^, N (%)** | Early (I–II) | 4 (7.5) | 0 (0) | 4 (10.8) |
|  | Advanced (III–IV) | 22 (41.5) | 47 (100) | 31 (83.8) |
|  | Unknown | 27 (51) | 0 (0) | 2 (5.4) |
| **Therapy, N (%)** | Surgery alone | 3 (5.7) | 2 (4.3) | 0 (0) |
|  | Surgery+Radiation | 2 (3.8) | 14 (29.8) | 0 (0) |
|  | Surgery+CRT | 4 (7.6) | 15 (31.9) | 0 (0) |
|  | Radiation alone | 2 (3.8) | 2 (4.3) | 0 (0) |
|  | CRT | 13 (24.4) | 13 (27.6) | 37 (100) |
|  | Incomplete data | 29 (54.7) | 1 (2.1) | 0 (0) |

^†^7^th^ edition AJCC staging manual

CRT=chemoradiotherapy; NA=Not Available

**Supplemental Table 2. Primers used for qPCR**

| **Gene** | **Forward** | **Reverse** |
| --- | --- | --- |
| **COX4I1** | CAGGGTATTTAGCCTAGTTGGC | GCCGATCCATATAAGCTGGGA |
| **MDH1** | TGCTGTCATCAAGGCTCGAA | CTCCCTCTGGGGTTCCAAAC |
| **ACADM** | GACTGAGGAGCCATTGATGTG | CCGTTGGTTATCCACATCTTCTG |
| **NDUFB8** | CTCCTTGTTGGGCTTATCACA | GCCCACTCTAGAGGAGCTGA |
| **SDHB** | AAGCATCCAATACCATGGGG | TCTATCGATGGGACCCAGAC |
| **UQCRC2** | GTTTGTTCATTAAAGCAGGCAGTAG | TGCTTCAATTCCACGGGTTATC |
| **ATP5A1** | ACTGGGCGTGTCTTAAGTATTG | ACCAAGGGCATCAACTACAC |
| **18S** | CTCAACACGGGAAACCTCAC | CGCTCCACCAACTAAGAACG |
| **MTCO1** | CCCACCGGCGTCAAAGTAT | TGCAGCAGATCATTTCATATTGC |
| **B2M** | TGCTGTCTCCATGTTTGATGTATCT | TCTCTGCTCCCCACCTCTAAGT |
| **HPV16E1** | AACGTGTTGCGATTGGTGTA | TACGCAATTTTGGAGGCTCT |
| **HPV16E2** | GCCAACACTGGCTGTATCAA | CATCCTGTTGGTGCAGTTAAA |
| **HPV16E4** | TCCAATGCCATGTAGACGAC | GCTCACACAAAGGACGGATT |
| **HPV16E5** | CCACAACATTACTGGCGTGC | GCAGAGGCTGCTGTTATCCAC |
| **HPV16E6** | TCAGGACCCACAGGAGCG | CCTCACGTCGCAGTAACTGTTG |
| **PPARGC1A** | CCAAGTCGTTCACATCTAGTTCA | TCTGAGTCTGTATGGAGTGACAT |
| **PPARGC1B** | CCACATCCTACCCAACATCAAG | CACAAGGCCGTTGACTTTTAGA |
| **ESRRA** | AGGGTTCCTCGGAGACAGAG | TCACAGGATGCCACACCATAG |

**Supplemental Table 3. Antibodies**

| **Target** | **Clonality** | **Immunogen** | **Isotype** | **Host** | **Dilution** | **Supplier** | **Catalog#** |
| --- | --- | --- | --- | --- | --- | --- | --- |
| **PGC1α** | Polyclonal | The C terminal region of human PGC-1α | IgG | Rabbit | 1:500 | Sigma Aldrich | HPA063136 |
| **PGC1α** | Polyclonal |  | IgG | Rabbit | 1:500 | Sigma Aldrich | SAB2106455 |
| **p53 (1C12)** | Monoclonal | Residues surrounding Ser20 of human p53 protein | IgG1 | Mouse | 1:1000 | Cell Signaling | 2524 |
| **p53 (7F5)** | Monoclonal | The amino terminus region of human p53 protein | IgG | Rabbit | 1:1000 | Cell Signaling | 2527 |
| **β-actin (AC-15)** | Monoclonal | The β-cytoplasmic N-terminal sequence | IgG1 | Mouse | 1:5000 | Sigma Aldrich | A3854 |
| **HA (C29F4)** | Monoclonal | HA-Tag | IgG | Rabbit | 1:500 | Cell Signaling | 3724 |
| **OxPhos Human Antibody**  Complex I-NDUFB8  Complex II-SDHB  Complex III-UQCRC2  Complex IV-COX II  Complex V- ATP5A | Cocktail, monoclonal | Full length protein | IgG | Mouse | 1:1000 | Thermo Fisher | 45-8199 |

**Supplemental Table 4. Plasmids**

| **Plasmid backbone** | **Insert** | **Promoter** | **Tag(s)** | **Selectable Marker** | **SOURCE** |
| --- | --- | --- | --- | --- | --- |
| pLentiN | none | CMV | Met/Flag/HA | Blasticidin | Addgene  #37444 |
| pLentiN 16E6no* | HPV16E6 (V42L) | CMV | Flag/HA | Blasticidin | Addgene  #37445 |
| pMD2.G | VSV G | CMV |  |  | Addgene  #12259 |
| psPAX2 | Gag-Pro-Pol |  |  |  | Addgene  #12260 |
| 2kB PGC-1α promoter | Human PGC-1α promoter | pGL3-basic | Luciferase |  | D. Kelly lab |
| MSCV-IP N FlagHA 16E6 | HPV16 E6 (V42L) | MSCV LTR | Flag/HA | Puromycin | E. White lab  Plasmid # 6724 |
| pMSCV-N-HAonly 16E6 | HPV16 E6 | MSCV LTR | HA | Puromycin | Addgene # 42603 |
| pMSCV-N-HA 16E6 8S9A10T | HPV16 E6 (8S9A10T) | MSCV LTR | HA | Puromycin | Addgene # 44153 |
| pMSCV-N-HA 16E6 I128T | HPV16 E6 (I128T) | MSCV LTR | HA | Puromycin | Addgene # 44154 |
| pMSCV-N-HA 16E6 Star | HPV16 E6 I* | MSCV LTR | HA | Puromycin | E. White lab  Plasmid # 7257 |
